## SupplementalFiles_AlvarezBuyllaetal for "Molecular physiology of pumiliotoxin sequestration in a poison frog": all_figures_plotting_and_statistics.R

```
## Aurora Alvarez-Buylla
## December 14, 2021
## Figure plotting and statistics scripts for manuscript entitled: 
## "Molecular physiology of pumiliotoxin sequestration in a poison frog"

#### ALL CODE SHOWN BELOW WAS RUN IN R VERSION 3.6.3 ######

#load libraries
library(readxl)		
library(ggplot2)	
library(reshape2)
library(ggpubr) 
library(RColorBrewer) 
library(dplyr)
library(car)
library(naniar)
library(patchwork)	
library(tidyverse)
library(glmmTMB)
library(FSA)
library(tidyr) 
library(reshape) 
library(lme4) 
library(emmeans)
library(broom) 

#################################################################################################################################################
############################# FIGURE 1 ##########################################################################################################
#################################################################################################################################################
# Alkaloid sequestration in different tissue types

##################### plotting #####################
tissue_order <- c("eggs","intestine","liver","skin")

#load excel file with data ("alkaloids" tab of the supplementary excel file)
LCMS_data <- read_excel("alkaloids.xlsx", .name_repair="universal")

#replace all "guts" with "intestine" label
LCMS_data <- LCMS_data %>% mutate(tissue = str_replace(tissue, "gut", "intestine"))

#turn tissue into factor
LCMS_data$tissue <- factor(LCMS_data$tissue , levels=tissue_order)

#replace "N/F" notation with NAs
LCMS_data_NAs <- LCMS_data %>% replace_with_na_at(.vars = c('aPTXc_norm_area'), condition = ~.x == "N/F")

#turn into numeric data
LCMS_data_NAs$aPTXc_norm_area <- as.numeric(LCMS_data_NAs$aPTXc_norm_area)

#replace NAs with 0s
LCMS_data_NAs$aPTXc_norm_area[is.na(LCMS_data_NAs$aPTXc_norm_area)] <- 0

#filter for just the skin
LCMS_data_NAs_skin <- LCMS_data_NAs[LCMS_data_NAs$tissue == "skin",]

#filter for only the ptx treated data
LCMS_data_NAs_ptx <- LCMS_data_NAs[LCMS_data_NAs$treatment == "DHQ+PTX",]

# plot DHQ pmols boxplot
dhq <- ggboxplot(LCMS_data, x="tissue", y="DHQ_pmol", fill="treatment") + 
  theme_bw() +
  theme(legend.position=c(0.2, 0.85), panel.grid.major = element_blank(), panel.grid.minor = element_blank(), panel.border = element_rect(size = 1, color="black"), plot.title = element_text(hjust = 0.5)) +
  ggtitle("Sequestered DHQ by tissue type") +
  xlab("tissue") +
  ylab("pmol of DHQ") +
  scale_fill_manual(values=c("darkgoldenrod1", "cornflowerblue")) + 
  geom_dotplot(aes(fill = treatment), binaxis='y', stackdir='center', dotsize = 0.7, position = position_dodge(0.8))

# plot PTX pmols boxplot
ptx <- ggboxplot(LCMS_data, x="tissue", y="PTX_pmol", fill="treatment") + 
  theme_bw() +
  theme(legend.position=c(0.2, 0.85), panel.grid.major = element_blank(), panel.grid.minor = element_blank(), panel.border = element_rect(size = 1, color="black"), plot.title = element_text(hjust = 0.5)) +
  ggtitle("Sequestered PTX by tissue type") +
  xlab("tissue") +
  ylab("pmol of PTX") +
  scale_fill_manual(values=c("darkgoldenrod1", "cornflowerblue")) + 
  geom_dotplot(aes(fill = treatment), binaxis='y', stackdir='center', dotsize = 0.7, position = position_dodge(0.8))

#plot the aPTX values in the DHQ+PTX treatment group
aPTXc <- ggboxplot(LCMS_data_NAs_ptx, x="tissue", y="aPTXc_norm_area", fill="treatment") + 
  theme_bw() +
  theme(legend.position=c(0.2, 0.89), panel.grid.major = element_blank(), panel.grid.minor = element_blank(), panel.border = element_rect(size = 1, color="black"), plot.title = element_text(hjust = 0.5))  +
  ggtitle("aPTX by tissue type") +
  xlab("tissue") +
  ylab("normalized aPTX AUC") +
  scale_fill_manual(values=c("cornflowerblue"))  +
  geom_dotplot(aes(fill = treatment), binaxis='y', stackdir='center', dotsize = 0.7, position = position_dodge(0.8))


# plot overall aPTX values in only the skin
aPTX_overall <- ggboxplot(LCMS_data_NAs_skin, x="treatment", y="aPTXc_norm_area", fill="treatment") + 
  theme_bw() +
  theme(legend.position=c(0.3, 0.85), panel.grid.major = element_blank(), panel.grid.minor = element_blank(), panel.border = element_rect(size = 1, color="black"), plot.title = element_text(hjust = 0.5))  +
  ggtitle("Skin aPTX") +
  xlab("treatment") +
  ylab("normalized aPTX AUC") +
  scale_fill_manual(values=c("darkgoldenrod1", "cornflowerblue"))  +
  geom_dotplot(aes(fill = treatment), binaxis='y', stackdir='center', dotsize = 0.7, position = position_dodge(0.8))

# patch the plots together
ptx_plots <- ptx + aPTX_overall + aPTXc + plot_layout(ncol=3,widths=c(1.5,1,1.5))

#save the plots
ggsave("dhq_plots.pdf", plot = dhq, width = 9.975, height = 10, units = c("cm"))
ggsave("ptx_plots.pdf", plot = ptx_plots, width = 26.6, height = 10, units = c("cm"))

# NOTE - figure aesthetics (color, label size) were minimally adjusted in Adobe illustrator

##################### statistics #####################
# glmm for DHQ pmol across tissues and treatment with frog as random effect
dhq_ml2 <- glmmTMB(DHQ_pmol ~ 1 + treatment * tissue + (1|frog),
        family="nbinom2", #negative binomial model
        ziformula=~1,  #zero inflation applying to all observations
        data=LCMS_data)
summary(dhq_ml2)
glmmTMB:::Anova.glmmTMB(dhq_ml2, type="II")

# glmm for PTX pmol across tissues and treatment with frog as random effect
ptx_ml2 <- glmmTMB(PTX_pmol ~ 1 + treatment * tissue + (1|frog),
        family="nbinom2", #negative binomial model
        ziformula=~1, #zero inflation applying to all observations
        data=LCMS_data)
summary(ptx_ml2)
glmmTMB:::Anova.glmmTMB(ptx_ml2, type="II")

#wilcoxon rank sum test to just look at aPTX differences in skin
just_skin <- wilcox.test(aPTXc_norm_area ~ group, data = LCMS_data_NAs_skin, 
	exact = FALSE)
just_skin

#Kruskall Wallis
just_aptx <- kruskal.test(aPTXc_norm_area ~ tissue, data = LCMS_data_NAs_ptx)
just_aptx

#Posthoc for K-W if significant: Dunn Test
just_aptx_ph = dunnTest(aPTXc_norm_area ~ tissue,
              data=LCMS_data_NAs_ptx,
              method="bh")   
just_aptx_ph

#################################################################################################################################################
############################# FIGURE 2 ##########################################################################################################
#################################################################################################################################################
# Differentially expressed genes in different tissues

##################### plotting #####################
#abundance values taken from TMM normalized values, can be found on "TMM_vals_genes_of_interest" tab of supplementary excel file

#gut genes of interest
abundances <- c(0,0,0,0,0,0,3.774,15.001,2.796,2.974)
treatment <- c("DHQ","DHQ","DHQ","DHQ","DHQ","DHQ+PTX","DHQ+PTX","DHQ+PTX","DHQ+PTX","DHQ+PTX")
CYP3A29 <- data.frame(treatment,abundances)
rownames(CYP3A29) <- c("AAB-E3-2","AAB-E3-10","AAB-E3-16","AAB-E3-36","AAB-E3-40","AAB-E3-6","AAB-E3-9","AAB-E3-20","AAB-E3-26","AAB-E3-39")

gut1 <- ggboxplot(CYP3A29, x="treatment", y="abundances", fill="treatment") + 
  theme_bw() +
  theme(legend.position="none", panel.grid.major = element_blank(), panel.grid.minor = element_blank(), panel.border = element_rect(size = 1, color="black"), plot.title = element_text(hjust = 0.5))  +
  ggtitle("CYP3A29") +
  xlab("") +
  ylab("TMM normalized counts") +
  scale_fill_manual(values=c("darkgoldenrod1", "cornflowerblue"))  +
  geom_dotplot(aes(fill = treatment), binaxis='y', stackdir='center', dotsize = 1, position = position_dodge(0.8))


abundances4 <- c(18.38,14.328,14.268,30.059,16.095,272.884,19.053,249.676,538.004,205.298)
MHCI <- data.frame(treatment, abundances4)
rownames(MHCI) <- c("AAB-E3-2","AAB-E3-10","AAB-E3-16","AAB-E3-36","AAB-E3-40","AAB-E3-6","AAB-E3-9","AAB-E3-20","AAB-E3-26","AAB-E3-39")
gut4 <- ggboxplot(MHCI, x="treatment", y="abundances4", fill="treatment") + 
  theme_bw() +
  theme(legend.position="none", panel.grid.major = element_blank(), panel.grid.minor = element_blank(), panel.border = element_rect(size = 1, color="black"), plot.title = element_text(hjust = 0.5), axis.title.y=element_blank())  +
  ggtitle(bquote("MHCI\u03b1")) +
  xlab("") +
  ylab("") +
  scale_fill_manual(values=c("darkgoldenrod1", "cornflowerblue"))  +
  geom_dotplot(aes(fill = treatment), binaxis='y', stackdir='center', dotsize = 1, position = position_dodge(0.8))

#liver genes of interest
abundances5 <- c(0,6.009,6.177,3.755,7.904,0,0,0,0,0)
VTG2 <- data.frame(treatment, abundances5)
rownames(VTG2) <- c("AAB-E3-2","AAB-E3-10","AAB-E3-16","AAB-E3-36","AAB-E3-40","AAB-E3-6","AAB-E3-9","AAB-E3-20","AAB-E3-26","AAB-E3-39")
liver1 <- ggboxplot(VTG2, x="treatment", y="abundances5", fill="treatment") + 
  theme_bw() +
  theme(legend.position="none", panel.grid.major = element_blank(), panel.grid.minor = element_blank(), panel.border = element_rect(size = 1, color="black"), plot.title = element_text(hjust = 0.5), axis.title.y=element_blank())  +
  ggtitle("VTG2") +
  xlab("") +
  ylab("") +
  scale_fill_manual(values=c("darkgoldenrod1", "cornflowerblue"))  +
  geom_dotplot(aes(fill = treatment), binaxis='y', stackdir='center', dotsize = 1, position = position_dodge(0.8))

abundances6 <- c(0,1.311,2.334,2.575,3.82,185.669,71.54,5.995,90.871,92.044)
MHCI2 <- data.frame(treatment, abundances6)
rownames(MHCI2) <- c("AAB-E3-2","AAB-E3-10","AAB-E3-16","AAB-E3-36","AAB-E3-40","AAB-E3-6","AAB-E3-9","AAB-E3-20","AAB-E3-26","AAB-E3-39")
liver2 <- ggboxplot(MHCI2, x="treatment", y="abundances6", fill="treatment") + 
  theme_bw() +
  theme(legend.position="none", panel.grid.major = element_blank(), panel.grid.minor = element_blank(), panel.border = element_rect(size = 1, color="black"), plot.title = element_text(hjust = 0.5), axis.title.y=element_blank())  +
  ggtitle(bquote("MHCI\u03b1")) +
  xlab("") +
  ylab("") +
  scale_fill_manual(values=c("darkgoldenrod1", "cornflowerblue"))  +
  geom_dotplot(aes(fill = treatment), binaxis='y', stackdir='center', dotsize = 1, position = position_dodge(0.8))

#skin genes of interest
abundances7 <- c(1.468,0.957,1.422,0.36,1.088,5.902,3.176,6.995,4.327,8.934)
treatment <- c("DHQ","DHQ","DHQ","DHQ","DHQ","DHQ+PTX","DHQ+PTX","DHQ+PTX","DHQ+PTX","DHQ+PTX")
STX1A <- data.frame(treatment,abundances)
rownames(STX1A) <- c("AAB-E3-2","AAB-E3-10","AAB-E3-16","AAB-E3-36","AAB-E3-40","AAB-E3-6","AAB-E3-9","AAB-E3-20","AAB-E3-26","AAB-E3-39")
skin1 <- ggboxplot(STX1A, x="treatment", y="abundances7", fill="treatment") + 
  theme_bw() +
  theme(legend.position="none", panel.grid.major = element_blank(), panel.grid.minor = element_blank(), panel.border = element_rect(size = 1, color="black"), plot.title = element_text(hjust = 0.5), axis.title.y=element_blank())  +
  ggtitle("STX1A") +
  xlab("") +
  ylab("") +
  scale_fill_manual(values=c("darkgoldenrod1", "cornflowerblue"))  +
  geom_dotplot(aes(fill = treatment), binaxis='y', stackdir='center', dotsize = 1, position = position_dodge(0.8))

abundances9 <- c(0,5.083,1.655,0.888,0.936,0,0,0,0,0)
SLC2 <- data.frame(treatment, abundances9)
rownames(SLC2) <- c("AAB-E3-2","AAB-E3-10","AAB-E3-16","AAB-E3-36","AAB-E3-40","AAB-E3-6","AAB-E3-9","AAB-E3-20","AAB-E3-26","AAB-E3-39")
skin3 <- ggboxplot(SLC2, x="treatment", y="abundances9", fill="treatment") + 
  theme_bw() +
  theme(legend.position="none", panel.grid.major = element_blank(), panel.grid.minor = element_blank(), panel.border = element_rect(size = 1, color="black"), plot.title = element_text(hjust = 0.5), axis.title.y=element_blank())  +
  ggtitle("SLC2") +
  xlab("") +
  scale_fill_manual(values=c("darkgoldenrod1", "cornflowerblue"))  +
  geom_dotplot(aes(fill = treatment), binaxis='y', stackdir='center', dotsize = 1, position = position_dodge(0.8))

#stitching plots together
gut1 | gut4 | liver1 | liver2 | skin1 | skin3

#saving plot
ggsave("goi.pdf", plot = last_plot(), width = 25, height = 8, units = c("cm"))

# NOTE - figure aesthetics (color, label size) were minimally adjusted in Adobe illustrator

##################### statistics #####################
## statistics taken from DEseq file and included in the supplementary excel file ##

#################################################################################################################################################
############################# FIGURE 3 ##########################################################################################################
#################################################################################################################################################
# Human CYP2D6 PTX 251D metabolism

##################### plotting #####################
# from "CYP_PTX_clearance" tab of supplementary excel file
df <- as.data.frame(read_excel("CYP_PTX_clearance.xlsx"))

# making Cheyenne's suggested line plot
cheyenne_plot <- ggplot(df, aes(x = incubation.time.minutes, y = percent.compound.remaining.MEAN, group = cyp)) +
   geom_line(aes(color = cyp), size = 1.3)+
   geom_point(aes(color = cyp), size = 3)+
   #geom_point(shape = 1,size = 3,colour = "black") +
   theme_bw()+
   theme(legend.title = element_blank(), panel.grid.major = element_blank(), panel.grid.minor = element_blank(), panel.border = element_rect(size = 1, color="black"), plot.title = element_text(hjust = 0.5)) +
   ggtitle("") + 
   xlab("time point (min)") +
   ylab("average % PTX 251D remaining") +
   scale_color_brewer(palette = "Dark2")

# making PTX metabolite bar plot  
# from "CYP2D6_PTX_metabolites" tab of supplementary excel file
df_aptx <- as.data.frame(read_excel("CYP2D6_PTX_metabolites.xlsx"))
df_aptx$time.point <- as.factor(df_aptx$time.point)
df_aptx$mz <- as.factor(df_aptx$mz)

colors <- c("#989898", "#FFFFFF") 
names(colors) <- c("0", "90")

paPTX <- ggplot(df_aptx, aes(fill=time.point, y=area, x=mz)) + 
  geom_bar(color="black", stat="identity") + 
  theme_bw() + 
  theme(legend.position=c(0.77, 0.85), panel.grid.major = element_blank(), panel.grid.minor = element_blank(), panel.border = element_rect(size = 1, color="black"), plot.title = element_text(hjust = 0.5)) +
  labs(fill = "time point (min)") + 
  ggtitle("Human CYP2D6 + PTX 251D metabolites") + 
  xlab("m/z (M+H)") +
  ylab("peak area") +
  scale_fill_manual(values=colors)

# stitching plots together
fig3plots <- cheyenne_plot + paPTX

#saving plot
ggsave("fig3plots.pdf", fig3plots, width = 25, height = 12, units = "cm")

# NOTE - figure aesthetics (color, label size) were minimally adjusted in Adobe illustrator

#################################################################################################################################################
############################# FIGURE 4 ##########################################################################################################
#################################################################################################################################################
# CYP2D6 expression and sequence conservation

##################### statistics #####################
#load the filtered TMM expression matrices for all tissues, found in tabs with corresponding name in supplementary excel file
gutTMM <- read.table(file = 'gut_CYP2D6_1e-115_blastHitMatches.gene.TMM.EXPR.matrix', sep = '\t', header = TRUE)
liverTMM <- read.table(file = 'liver_CYP2D6_1e-115_blastHitMatches.gene.TMM.EXPR.matrix', sep = '\t', header = TRUE)
skinTMM <- read.table(file = 'skin_CYP2D6_1e-115_blastHitMatches.gene.TMM.EXPR.matrix', sep = '\t', header = TRUE)

#convert the TMM matrices to long format
gutTMM <- melt(gutTMM, id=c("X"))
liverTMM <- melt(liverTMM, id=c("X"))
skinTMM <- melt(skinTMM, id=c("X"))

#load the sample data for all tissues
samples <- read.table(file = 'samples.txt', sep = '\t', header = TRUE)

#merge sample data with each TMM matrix
gutMerged <- merge(gutTMM, samples, by='variable')
liverMerged <- merge(liverTMM, samples, by='variable')
skinMerged <- merge(skinTMM, samples, by='variable')

#rename and remove unecessary columns for gut
gutMerged <- dplyr::rename(gutMerged, 'sample' = 'variable')
gutMerged <- dplyr::rename(gutMerged, 'contig' = 'X')
gutMerged <- dplyr::rename(gutMerged, 'tmm.expression' = 'value')
# gutMerged <- dplyr::rename(gutMerged, 'treatment' = 'V2')
# gutMerged <- dplyr::rename(gutMerged, 'tissue' = 'V3')
gutMerged$sample <- NULL
gutMerged$tissue <- NULL

#rename and remove unecessary columns for liver
liverMerged <- dplyr::rename(liverMerged, 'sample' = 'variable')
liverMerged <- dplyr::rename(liverMerged, 'contig' = 'X')
liverMerged <- dplyr::rename(liverMerged, 'tmm.expression' = 'value')
# liverMerged <- dplyr::rename(liverMerged, 'treatment' = 'V2')
# liverMerged <- dplyr::rename(liverMerged, 'tissue' = 'V3')
liverMerged$sample <- NULL
liverMerged$tissue <- NULL

#rename and remove unecessary columns for skin
skinMerged <- dplyr::rename(skinMerged, 'sample' = 'variable')
skinMerged <- dplyr::rename(skinMerged, 'contig' = 'X')
skinMerged <- dplyr::rename(skinMerged, 'tmm.expression' = 'value')
# skinMerged <- dplyr::rename(skinMerged, 'treatment' = 'V2')
# skinMerged <- dplyr::rename(skinMerged, 'tissue' = 'V3')
skinMerged$sample <- NULL
skinMerged$tissue <- NULL

#turn treatment into a factor
gutMerged$treatment <- as.factor(gutMerged$treatment)
liverMerged$treatment <- as.factor(liverMerged$treatment)
skinMerged$treatment <- as.factor(skinMerged$treatment)

#run the t-test for gut
gutResults <- gutMerged %>%
  group_by(contig) %>%
  do(tidy(t.test(.$tmm.expression ~ .$treatment)))

#benjamini hochberg correct p-values for multiple hypothesis testing
gutResults$p.adjusted <- p.adjust(gutResults$p.value, method = "BH")
#filter(gutResults, p.value <= 0.05)$contig
#filter(gutResults, p.adjusted <= 0.05)

#run the t-test for liver
liverResults <- liverMerged %>%
  group_by(contig) %>%
  do(tidy(t.test(.$tmm.expression ~ .$treatment)))

#benjamini hochberg correct p-values for multiple hypothesis testing
liverResults$p.adjusted <- p.adjust(liverResults$p.value, method = "BH")
#filter(liverResults, p.value <= 0.05)$contig
#filter(liverResults, p.adjusted <= 0.05)

#run the t-test for skin
skinResults <- skinMerged %>%
  group_by(contig) %>%
  do(tidy(t.test(.$tmm.expression ~ .$treatment)))

#benjamini hochberg correct p-values for multiple hypothesis testing
skinResults$p.adjusted <- p.adjust(skinResults$p.value, method = "BH")
#filter(skinResults, p.value <= 0.05)$contig
#filter(liverResults, p.adjusted <= 0.05)

#filter data for the only significant hit (TRINITY_DN15846_c0_g2 in the gut)
gutDN15846 <- gutMerged[gutMerged$contig == "TRINITY_DN15846_c0_g2",]

#run a simple t-test between treatments to get the p-value
t.test(gutDN15846$tmm.expression[gutDN15846$treatment=="DHQ"], gutDN15846$tmm.expression[gutDN15846$treatment=="DHQ_PTX"])

##################### plotting #####################

pDN15846 <- ggplot(gutDN15846, aes(x=treatment, y=tmm.expression, fill=treatment)) + 
  theme_bw() +
  theme(legend.position="none", 
  	panel.grid.major = element_blank(), 
  	panel.grid.minor = element_blank(), 
  	panel.border = element_rect(size = 1, color="black"), 
  	plot.title = element_text(hjust = 0.5))  +
  ggtitle("TRINITY_DN15846_c0_g2") +
  xlab("treatment") +
  ylab("TMM normalized counts") +
  geom_boxplot(outlier.shape = NA) +
  scale_fill_manual(values=c("darkgoldenrod1", "cornflowerblue"))  +
  geom_dotplot(aes(fill = treatment), binaxis='y', stackdir='center', dotsize = 1, position = position_dodge(0.8))

ggsave("TRINITY_DN15846_c0_g2.pdf", plot = pDN15846, width = 8, height = 10, units = c("cm"))

# NOTE - figure aesthetics (color, label size) were minimally adjusted in Adobe illustrator

# Panel 4B was made with blast+ command line tools, Benchling, and Adobe illustrator as detailed in the methods
```
