## SupplementalFiles_AlvarezBuyllaetal for "Molecular physiology of pumiliotoxin sequestration in a poison frog": binSeqsByLength.pl

```
#!/oak/stanford/groups/loconnel/miniconda/bin/perl 

=head1 Name

binSeqsByLength.perl

=head1 Usage

binSeqsByLength.perl [-s] [-b "100 200 ..."] inputFastaFile [inputFastaFile ...]

=head1 Synopsis

Takes a fasta input file or files, bins the sequences by length (0 to 99, 100 to 199, etc) and writes the results
to multiple FASTA files.

Options: -s  = sort by size within bins
         -b <binning boundaries> = use alternate bin sizes
	 -f = force overwriting of existing files

=head1 Notes

This script asssumes you have provided a fasta format input file. It does not do any error checking for this. 

You will need Bioperl installed, in particular, the SeqIO.pm module.

=head1 Author

Tim Booth, NEBC

=cut

#start code
our $VERSION = 1.0;

use strict;
use warnings;
# use Data::Dumper;
use Bio::SeqIO;
use Getopt::Std;

our ($opt_s, $opt_b, $opt_h, $opt_f);
getopts('hfsb:');

if ($opt_h) {die "Usage: binSeqsByLength.perl [-s] [-b \"100 200 ...\"] inputFastaFile [inputFastaFile ...] \n";}

my $infileFormat = 'fasta';
my $outfileFormat = 'fasta';

my $infileName;
if(@ARGV == 0 || $ARGV[0] eq '-') { $infileName = "input_data.$infileFormat"; }
elsif(@ARGV == 1)                 { ($infileName) = $ARGV[0] =~ /(?:.*\/|^)(.*)/; }
else                              { $infileName = "input_files.$infileFormat"; }

my $seq_stream = Bio::SeqIO->newFh(-format => $infileFormat,
                                   -fh     => \*ARGV);

# loads the whole file into memory - be careful
# if this is a big file, then this script will
# use a lot of memory
our ($count, $writecount) = (0, 0);
our (@bins, @bin_tops, $bin_max, $last_bin_repeat, $last_repeat_width, $last_bin_idx);

#Now analyse the -b option if there was one.
if(defined($opt_b))
{
    $last_bin_repeat = ($opt_b =~ /\.\.\.$/); #See if the list ends '...'
    @bin_tops = sort {$a <=> $b} grep {$_ && ! /[^0-9]/} split /\s+/, $opt_b;

    #In repeat mode we need at least two bins
    if($last_bin_repeat && scalar(@bin_tops) < 2) { die "Invalid boundaries - see instructions.\n"; }
    
    $bin_max = $bin_tops[-1] || 0;
    $last_bin_idx = scalar(@bin_tops);
    if($last_bin_repeat)
    {
	$last_repeat_width = $bin_tops[-1] - $bin_tops[-2];
    }
}
else
{
    #Default is "100 200 ..."
    @bin_tops = (100);
    $bin_max = 0;
    $last_bin_repeat = 1;
    $last_repeat_width = 100;
    $last_bin_idx = 0;
}    

#Now some weird logic to figure out what bin to put the sequence in:
sub length_to_bin_idx
{
    my ($length) = @_;
    my $bin = 0;
    if($length < $bin_max)
    {
	for(@bin_tops)
	{
	    last if ($length < $_);
	    $bin++;
	}
    }
    else
    {
	if($last_bin_repeat)
	{
	    $bin = int(($length - $bin_max) / $last_repeat_width) + $last_bin_idx;
	}
	else
	{
	    $bin = $last_bin_idx;
	}
    }
    $bin;
}

#Now the main code
#Read 'em
while ( my $seq = <$seq_stream> ) 
{
    
    my $bin = length_to_bin_idx($seq->length);
    {
	push @{$bins[$bin]}, $seq;
    }
    $count ++;
}

#Dump 'em
for(my $nn = 0; $nn < @bins; $nn++)
{
    my $abin = $bins[$nn];

    #Skip empties
    next unless defined($abin) && scalar(@$abin);
    my $binsize = scalar(@$abin);

    #Make a name
    my $name = ''; my $outfileName;
    if(!@bin_tops) #Useful if you only wanted to sort.
    {
	$name = 'all';
    }
    elsif($nn == 0)
    {
	$name = "0_to_" . ($bin_tops[$nn] - 1);
    }
    elsif($nn < @bin_tops)
    {
	$name = $bin_tops[$nn-1] . "_to_" . ($bin_tops[$nn] - 1);
    }
    elsif($last_bin_repeat)
    {
	$name = ((($nn - $last_bin_idx) * $last_repeat_width) + $bin_max) . "_to_" .
	        ((($nn + 1 - $last_bin_idx) * $last_repeat_width) + $bin_max - 1);
    }
    else
    {
	$name = $bin_max . "_or_more";
    }

    if($infileName =~ /(.*?)\.?$infileFormat/i)
    {
	$outfileName = "${1}_$name.$outfileFormat";
    }
    else
    {
	$outfileName = "${infileName}_$name";
    }

    print "Writing $binsize sequences to $outfileName\n";

    if(-e $outfileName)
    {
	if($opt_f) { unlink($outfileName) || die "** Could not remove old file - aborting.\n" }
	else	   { die "** File already exists - aborting.\n" }
    }
    
    #Open output
    my $seq_out = Bio::SeqIO->new('-file' => ">$outfileName",
				  '-format' => $outfileFormat);

    #Dump seqs, maybe with sorting
    if($opt_s)
    {
	$seq_out->write_seq($_) for sort {$a->length <=> $b->length} @$abin;
    }
    else
    {
	$seq_out->write_seq($_) for @$abin;
    }

    $writecount++;
}

#print a message to tell the world of my awesomeness.
print "Done - $count sequences split into $writecount files". ($opt_s ? ' and sorted' : '') . "\n";
```
