## SupplementalFiles_AlvarezBuyllaetal for "Molecular physiology of pumiliotoxin sequestration in a poison frog": get_notselected_sequences.pl

```
#!/oak/stanford/groups/loconnel/miniconda/bin/perl -w
#############################
###   Jennifer Meneghin   ###
###   February 9, 2011    ###
#############################
#---------------------------------------------------------------------------------------------------------------------------
#Deal with passed parameters
#---------------------------------------------------------------------------------------------------------------------------
if ($#ARGV == -1) {
    &usage;
}
$in_file = "";
$fasta_file = "";
$out_file = "NotSelectedSequences.fa";
%my_args = @ARGV;
for $i (sort keys %my_args) {
    if ($i eq "-i") {
	$in_file = $my_args{$i};
    }
    elsif ($i eq "-f") {
	$fasta_file = $my_args{$i};
    }
    elsif ($i eq "-o") {
	$out_file = $my_args{$i};
    }
    else {
	print "\nUnrecognized argument: $i\n\n";
	&usage;
    }
}
unless ( open(IN, "$in_file") ) {
    print "\nGot a bad sequence ID list file: $in_file\n\n";
    &usage;
}
unless ( open(FASTA, "$fasta_file") ) {
    print "\nGot a bad FASTA file: $fasta_file\n\n";
    &usage;
}
unless ( open(OUT, ">$out_file") ) {
    print "\nGot a bad output file: $out_file\n\n";
    &usage;
}
print "Parameters:\nsequence ID list file = $in_file\nFASTA file = $fasta_file\noutput file = $out_file\n\n";
#---------------------------------------------------------------------------------------------------------------------------
#The main event
#---------------------------------------------------------------------------------------------------------------------------
%records = ();
while (<IN>) {
    chomp;
    s/\r//g;
    $records{$_} = $_;
}
$seq = "";
$header = "";
while (<FASTA>) {
    if (/^>/) {
	if (length($header) > 0) {
	    if (!$records{$id}) {
		print ">$header\n";
		print OUT ">$header\n";
		print OUT "$seq";
	    }
	    $header = "";
	    $seq = "";
	}
	$header = $_;
	$header =~ s/>//g;
	chomp($header);
	$id = $header;
	$id =~ s/^(.+?)\s.*$/$1/g;
    }
    else {
	$seq = $seq . $_;
    }
}
if (length($header) > 0) {
    if (!$records{$id}) {
	print ">$header\n";
	print OUT ">$header\n";
	print OUT "$seq";
    }
}
close(IN);
close(FASTA);
close(OUT);
#-----------------------------------------------------------------------
sub usage {
    print "\nUsage: ./get_notselected_sequences.pl\n\n";
    print "Parameters:\n";
    print "-i input file\tA Sequence ID List file that contains one sequence ID per line.\n";
    print "-f input file\tA FASTA file.\n";
    print "-o output file\tReturns a fasta file with only the sequences from the original FASTA sequence file\n";
    print "\t\tthat are NOT in the original file.\n\n";
    print "This script selects the sequences from the original fasta file that are NOT in the sequence id list file.\n\n";
    print "Jennifer Meneghin\n";
    print "February 9, 2011\n\n";
    print "Updated December 12, 2012\n\n";
    exit;
}
#-----------------------------------------------------------------------
```
