## SupplementalFiles_AlvarezBuyllaetal for "Molecular physiology of pumiliotoxin sequestration in a poison frog": RNAseq_analysis_scripts.pl

```
## Aurora Alvarez-Buylla
## December 14, 2021
## RNAseq analysis information for manuscript entitled: 
## "Molecular physiology of pumiliotoxin sequestration in a poison frog"

## Information and reccomendations for this pipeline were taken from
## https://github.com/trinityrnaseq/trinityrnaseq/wiki/Running-Trinity

## Associated files included in supplementary material: 

## dtf_samples_file.txt 
## contaminants_2019.txt
## get_notselected_sequences.pl
## binSeqsByLength.pl
## skin_samples.txt 
## liver_samples.txt 
## gut_samples.txt 

##################################################
##### Step 1: Assemble de-novo transcriptome #####
##################################################

#first, activate miniconda (path is user specific)
source activate $PATH/miniconda

# first this was attempted and ran out of memory
Trinity \
	--seqType fq \
	--SS_lib_type RF \
	--samples_file dtf_samples_file.txt \
	--output trinity \
	--max_memory 100G \
	--CPU 16 \
	--trimmomatic
	--full_cleanup

TrinityStats.pl trinity.Trinity.fasta

# then the normalized reads from the previous run were used
# in the new attempt and the normalization was skipped
Trinity \
	--seqType fq \
	--SS_lib_type RF \
	--left left.norm.fq \
	--right right.norm.fq \
	--output trinity \
	--max_memory 1000G \
	--trimmomatic \
	--CPU 20 \
	--full_cleanup \
	--no_normalize_reads

TrinityStats.pl trinity.Trinity.fasta

##########################################################
##### Step 2: Annotating and filtering transcriptome #####
##########################################################

# annotating the transcriptome using Swiss-Prot
DB="PATH/swissprot" 				# location of swiss-prot database
FA="trinity.Trinity.fasta" 			# assembled de-novo transcriptome
OUT=${FA//.fasta/.blastx.outfmt6}	# name for output file

#blastx command
blastx -query $FA -db $DB -num_threads 8 -max_target_seqs 1 -evalue 1e-10 -outfmt 6 > $OUT


#### filtering out contaminant contigs (i.e. microbial) ####
export DB=$PATH/db 		#path to list of contaminant organisms
export BIN=$PATH/bin 	#path to perl scripts used (see associated files)

# Get the names of the genes from the blast file, split apart the gene name file, and select the list or organisms from the split file. 
perl -e ' @cols=(1); while(<>) { s/\r?\n//; @F=split /\t/, $_; print join("\t", @F[@cols]), "\n" } warn "\nChose columns ", join(", ", @cols), " for $. lines\n\n" ' trinity.Trinity.blastx.outfmt6 > genename.txt

perl -e ' $sep="_"; while(<>) { s/\Q$sep\E/\t/g; print $_; } warn "Changed $sep to tab on $. lines\n" ' genename.txt > genenamenotab.txt

perl -e ' @cols=(1); while(<>) { s/\r?\n//; @F=split /\t/, $_; print join("\t", @F[@cols]), "\n" } warn "\nChose columns ", join(", ", @cols), " for $. lines\n\n" ' genenamenotab.txt > organismsinblast.txt

#compare your blast hits to the contaminant list to find overlap 
perl -e ' ($file1, $file2) = @ARGV; $printed = 0; open F2, $file2; while (<F2>) { $h2{$_}++ }; $count2 = $.; open F1, $file1; while (<F1>) { if (! $h2{$_}) { print $_; $printed++; } } $count1 = $.; warn "\nRead $count1 lines from $file1 and $count2 lines from $file2.\nPrinted $printed lines found in $file1 but not in $file2\n\n" ' organismsinblast.txt $DB/contaminants_2019.txt > postfilter.txt  

## filter out blast hits present in the contaminants list
# get contigs with contaminant hits
grep -f $DB/contaminants_2019.txt trinity.Trinity.blastx.outfmt6 > contamblast.txt

# select the contig names from the above contamblast.txt list
perl -e ' @cols=(0); while(<>) { s/\r?\n//; @F=split /\t/, $_; print join("\t", @F[@cols]), "\n" } warn "\nChose columns ", join(", ", @cols), " for $. lines\n\n" ' contamblast.txt > contamcontigs.txt

# select sequences in fasta file not in the contamcontigs.txt file
perl $BIN/get_notselected_sequences.pl -i contamcontigs.txt –f trinity.Trinity.fasta -o trinity.Trinity.cleanedcontigs.fasta

##################################################
##### Step 3: Clustering overlapping contigs #####
##################################################
# using cd-hit-est to cluster overlapping contigs

#first, activate miniconda (path is user specific)
source activate $PATH/miniconda

FA="trinity.Trinity.cleanedcontigs.fasta"
OUT=${FA//.fasta/_cdhit98.fasta}

cd-hit-est -T 24 -M 128000 -i $FA -o $OUT -c 0.98 -n 8 

########################################
##### Step 4: Remove small contigs #####
########################################
# remove contigs shorter than 250 bp

#first, activate miniconda (path is user specific)
source activate $PATH/miniconda

$PATH/bin/binSeqsByLength.pl -b '250' trinity.Trinity.cleanedcontigs_cdhit98.fasta 

# output (trinity_cdhit98_250_or_more.fasta) renamed after this step using the command:
mv trinity_cdhit98_250_or_more.fasta rinity.Trinity.cleanedcontigs.longreads.fasta

################################################################
##### Step 5: Prepare reference transcriptome for assembly #####
################################################################

#first, activate miniconda (path is user specific)
source activate $PATH/miniconda

FA="trinity.Trinity.cleanedcontigs.longreads.fasta"

# this perl script is part of the trinity toolkit
# https://github.com/trinityrnaseq/trinityrnaseq/wiki/Trinity-Transcript-Quantification
align_and_estimate_abundance.pl \
	--transcripts $FA \
	--est_method kallisto \
	--output_dir index \
	--trinity_mode \
	--prep_reference

#################################################################
##### Step 6: Align reads and calculate abundance estimates #####
#################################################################
# this loop was used to call the alignKallisto.sbatch script below
# passing in the paired quality trimmed reads for alignment and abundance estimates
# LOOP:
for i in ../trinity/AAB-E3-*_R1_001.fastq.gz.P.qtrim.gz
do 
  sbatch alignKallisto.sbatch $i ${i/_R1_001.fastq.gz.P.qtrim.gz/_R2_001.fastq.gz.P.qtrim.gz}
done

# the below code was part of an sbatch script called by the above loop
# SBATCH script: 
########################## alignKallisto.sbatch ##########################
#first, activate miniconda (path is user specific)
source activate $PATH/miniconda

R1=$1
R2=$2
FA="trinity.Trinity.cleanedcontigs.longreads.fasta"
sample=$(basename ${R1%%_R1*})

# this perl script is part of the trinity toolkit
# https://github.com/trinityrnaseq/trinityrnaseq/wiki/Trinity-Transcript-Quantification
align_and_estimate_abundance.pl \
  --transcripts $FA \
  --seqType fq \
  --SS_lib_type RF \
  --left $R1 \
  --right $R2 \
  --est_method kallisto \
  --thread_count 8 \
  --trinity_mode \
  --output_dir $sample
#############################################################################

#################################################################
##### Step 7: Split analysis into different groups (tissue) #####
#################################################################

# from this point forward analysis proceeds independently + in 
# parallel for each tissue type

# separate directories were made for each tissue type and individual sbatch
# scripts for each of the following steps were used. A generic "tissue" label
# will be used in the following steps 

mkdir gut
mkdir liver
mkdir skin

######################################################
##### Step 8: Make matrix of abundance estimates #####
######################################################
#first, activate miniconda (path is user specific)
source activate $PATH/miniconda

# this perl script is part of the trinity toolkit
# https://github.com/trinityrnaseq/trinityrnaseq/wiki/Trinity-Transcript-Quantification
# pass in the abundance estimates for all of the files associated with that tissue
# in this example the files associated with the skin are passed. 
abundance_estimates_to_matrix.pl \
--est_method kallisto \
--gene_trans_map ../trinity.Trinity.cleanedcontigs.longreads.fasta.gene_trans_map \
--out_prefix tissue.kallisto \
--name_sample_by_basedir ../AAB-E3-3/abundance.tsv ../AAB-E3-12/abundance.tsv ../AAB-E3-14/abundance.tsv ../AAB-E3-17/abundance.tsv ../AAB-E3-25/abundance.tsv ../AAB-E3-13/abundance.tsv ../AAB-E3-19/abundance.tsv ../AAB-E3-21/abundance.tsv ../AAB-E3-22/abundance.tsv ../AAB-E3-23/abundance.tsv

#############################################################
##### Step 9: Find differentially expressed transcripts #####
#############################################################
source activate $PATH/miniconda

# this perl script is part of the trinity toolkit
# https://github.com/trinityrnaseq/trinityrnaseq/wiki/Trinity-Differential-Expression
run_DE_analysis.pl \
	--matrix tissue.kallisto.gene.counts.matrix \
	--method DESeq2 \
	--samples_file tissue_samples.txt \
	--output tissue.kallisto.gene.counts.matrix

#################################################################
##### Step 10: Analyze differentially expressed transcripts #####
#################################################################

# this perl script is part of the trinity toolkit
# https://github.com/trinityrnaseq/trinityrnaseq/wiki/Trinity-Differential-Expression
# p-value cutoff of 0.5 and 4-fold change
analyze_diff_expr.pl --matrix ../tissue.kallisto.gene.TMM.EXPR.matrix --samples ../tissue_samples.txt -P 0.05 -C 2
```
